# Ancestral genome reconstruction enhances transposable element annotation by identifying degenerate integrants

**DOI:** 10.1101/2023.04.30.537485

**Authors:** Wayo Matsushima, Evarist Planet, Didier Trono

## Abstract

Growing evidence indicates that transposable elements (TEs) play important roles in evolution by providing genomes with coding and non-coding elements. Identification of TE-derived functional elements, however, has relied on TE annotations in individual species, which limits its scope to relatively intact TE sequences and misses elements derived from evolutionarily old TEs. Here, we report a novel approach to uncover previously unannotated degenerate TEs (degTEs) by probing multiple ancestral genomes reconstructed from hundreds of species. We applied this method to the human genome and discovered 1,452,810 degTEs, representing a 10.8% increase over the most recent human TE coverage. Further, we discovered that degTEs contribute to various *cis*-regulatory elements as well as transcription factor binding sites, including those of a known TE-controlling family, the KRAB zinc-finger proteins. We also report unannotated chimeric transcripts between degTEs and human genes expressed in embryos. This study provides a novel methodology and a freely available resource that will facilitate the investigation of TE co-option events on a full scale.

## Introduction

Transposable elements (TEs) are genetic units capable of mobilising their sequence within the host genome. In the most recent telomere-to-telomere (T2T) complete human genome assembly, 54% of the genomic DNA is estimated to be derived from TEs and repetitive elements, consisting mostly of long terminal repeats (LTRs), long and short interspersed nuclear elements (LINEs and SINEs, respectively) and DNA transposons^1^. However, the vast majority of the sequences in the human genome annotated as TE are transposition-incompetent due to mutations or insertions/deletions (indels) accumulated during evolution, and only an estimated 1:1,000 integrants belonging to the L1, Alu, SVA or HERV-K families remain active^2^.

It has become apparent that transposition-incompetent TEs are not just fossils of parasitic sequences. Indeed, TEs are frequently co-opted as regulatory elements (reviewed in ref^3^), and, at times, even give rise to new genes (reviewed in refs^4,5^). TEs are also spliced into genic transcripts, resulting in TE-gene chimeric transcripts, or transpochimeric gene transcripts (TcGTs), often produced in specific cell types as alternatives to canonical transcripts^6,7^. Since each lineage has acquired distinct TE subfamilies during evolution, TEs contribute to lineage-specific genomic innovations.

Currently, the gold-standard method to annotate TEs in an assembled genome is repository-based annotation, best exemplified by RepeatMasker^8^. This software scans a genome and identifies loci that significantly resemble one of the consensus sequences registered on a TE repository such as Repbase^9^ (https://www.girinst.org/repbase/) or Dfam^10^ (https://www.dfam.org/). Thus, this methodology is inherently limited to discovery of TEs that still retain a high sequence similarity to the consensus and might miss highly degenerate TEs, typically evolutionarily old elements.

Recent efforts have provided whole-genome sequences from an increasing number of species, including more than 600 vertebrates^11,12^. Armstrong et al. described a scalable method to align large numbers of genomes through the reconstruction of their ancestral sequences^13^. These reconstructed ancestral genomes (RAGs) are the best proxy to the genomes of extinct common ancestors, which are otherwise challenging to obtain because of difficulties in tracing back lineages and assembling sequences obtained from fossilised materials. Since ancestral genomes are reconstructed correcting for later-occurring mutational changes based on phylogeny, the sequences of their TEs are, in theory, closer to the consensus. This was in part confirmed for the L1PA6 subfamily^13^ but has not been verified for other TEs.

Here, we present a novel method that utilises multiple RAGs to identify TEs that escape common annotation techniques. It allowed us to discover previously unannotated integrants belonging to all the major TE classes in the human genome, extending its total TE coverage by 10.8%. We further found these newly unearthed TEs to contribute various *cis*-regulatory elements as well as transcription factor binding sites and to form chimeric transcripts with human genes expressed at specific embryonic stages.

## Results

### TE annotation in RAGs

We present a versatile method to identify degenerate TEs using multiple reconstructed ancestral genomes (RAGs) (Fig. 1). While TE integrants detected in older ancestral genomes are expected to be closer to the consensus because of a shorter divergence time since the initial emergence of the corresponding TE subfamilies, RAGs are devoid of TE subfamilies that appeared later in evolution (e.g. the primate ancestral genome does not harbour human-specific TEs). Thus, to recover TEs that emerged at wide-ranged evolutionary timepoints, multiple RAGs ancestral to a species of interest (SOI) are used.

**Figure 1.**
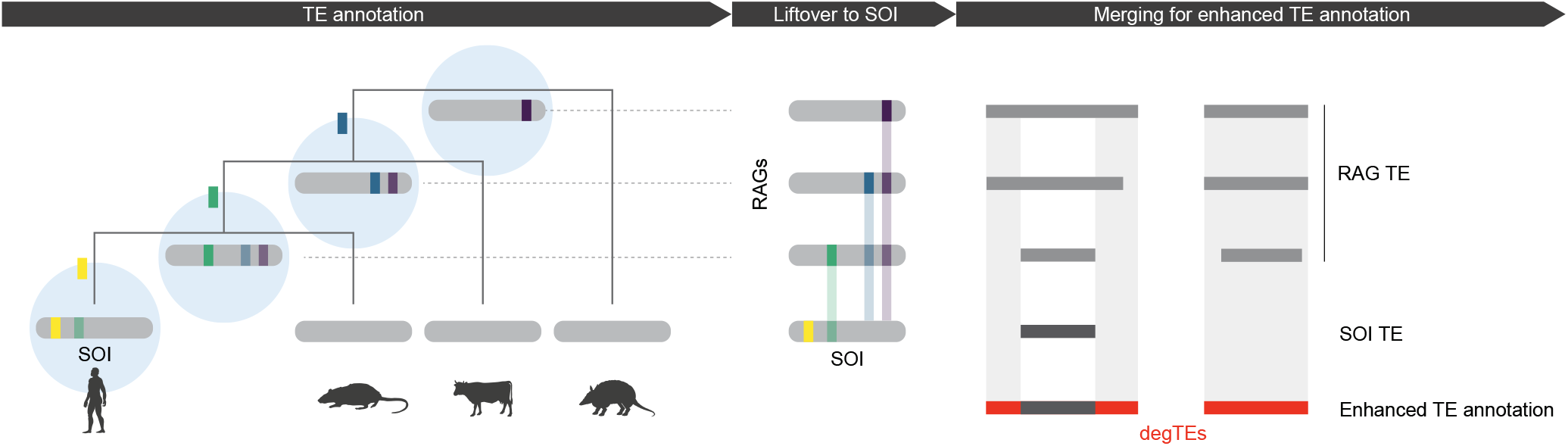
Overview of the RAG-enhanced TE annotation. Schematic representation of the workflow to achieve enhanced TE annotation by utilising multiple RAGs. Firstly, TEs are annotated in the species of interest (SOI) as well as in RAGs (blue circles). TEs annotated in RAGs are then lifted over to corresponding regions in SOI. Finally, lifted-over TE annotations not overlapping TEs in SOI (degTEs) are either merged to extend existing annotations (left) or added as new integrants (right). TEs inserted at different evolutionary timepoints are shown as coloured boxes with transparency indicating their divergence level from the consensus.

First, TEs are annotated in the RAGs and in the SOI. The RAG TEs are then lifted over to the SOI genome to find their corresponding sites. Finally, the lifted-over TEs are compared against the SOI TEs. If they do not overlap, the lifted-over TEs are added as novel elements, and the ones with a partial overlap are merged to extend existing integrants in the SOI genome.

As a proof of principle, we enhanced TE annotation in the human genome assembly GRCh38 (hg38), using six RAGs corresponding respectively to the common ancestors of Simiiformes, primates, Euarchonta, Euarchontoglires, Boreoeutheria, and Eutheria (Fig. 2a). Employing the same RepeatMasker parameters, we annotated TEs in hg38 (hereafter, hg38 TEs) and in RAGs, 30-50% of which were attributed to TEs (Fig. 2b). We found that the vast majority of the TEs identified in each RAG were older than their host genome as anticipated (Fig. 2c). Furthermore, TE divergence profiles revealed that a peak corresponding to evolutionarily young LINE-1, Alu, and SVA elements was only seen in hg38 and the simian RAG and was absent in the older RAGs (Fig. 2d). Together, these results confirm that precise reconstruction of TEs was achieved in the RAGs.

**Figure 2.**
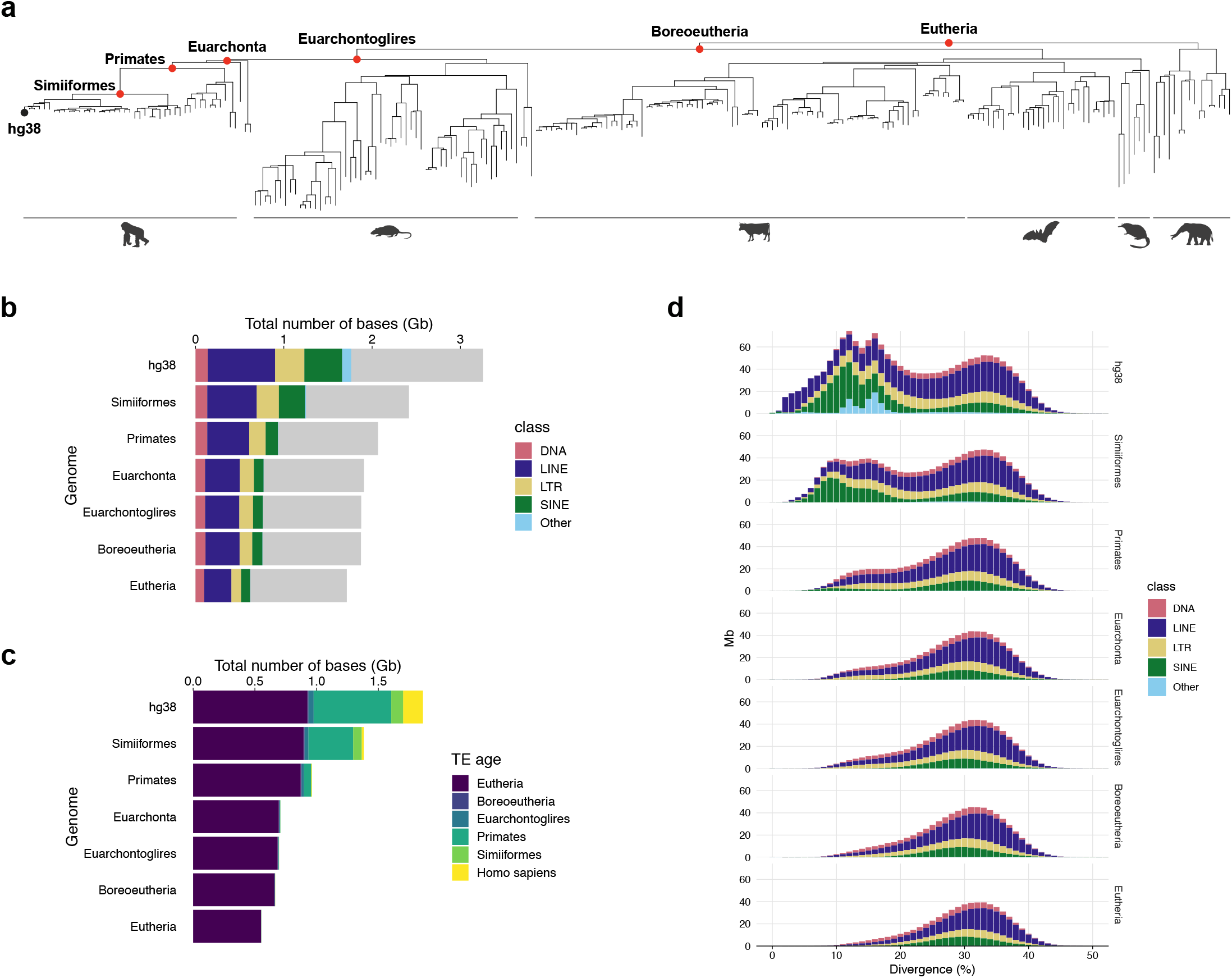
Characterisations of TEs annotated in the RAGs. (a) A phylogenetic tree of 241 species used to reconstruct the ancestral genomes (red points) for enhanced hg38 TE annotation. (b) Coverage of TE classes detected in each RAG and in hg38. Grey bars represent the non-TE regions in each genome. (c) TE age proportions in hg38 and the RAGs. TEs born outside of these six age categories were classified into the next younger age category (e.g. TEs born between primates and Simiiformes common ancestors were classified as Simiiformes). (d) Distribution of divergence levels of TE integrants in hg38 and the RAGs. The divergence level was rounded to a nearest integer.

### RAGs enhanced human TE annotation by unbiased identification of degTEs

For the TE annotation of each RAG, corresponding genomic loci in hg38 were sought using halLiftover^14^ (Supplementary Fig. 1a). Of the TEs found in each RAG, between 60-130 Mb corresponded to genomic sequences unassigned to TEs in hg38, which, when combined, amounted to 191 Mb of additional sequences annotated as TEs (Supplementary Fig. 1b).

We will refer to such newly identified TE-derived elements in hg38 as degenerate TEs (degTEs) for simplicity (Fig. 1). As shown in Fig. 3a, degTEs either represent newly discovered integrants (MamSINE1, MamRTE1, and L2a) or extend already annotated inserts (L2d). Also, distinct integrants were discovered in different RAGs. MamRTE1 integrants were detected in a relatively young RAG, the primate common ancestor, while L2a inserts were identified only when going up to the Euarchontoglires common ancestor, presumably because genetic drift erased their signature features after that. Conversely, the primate-specific AluSg integrant was not annotated in the RAGs older than the simian common ancestor, which suggests that it inserted in the middle of the MLT1 element between the primate and simian common ancestors.

**Figure 3.**
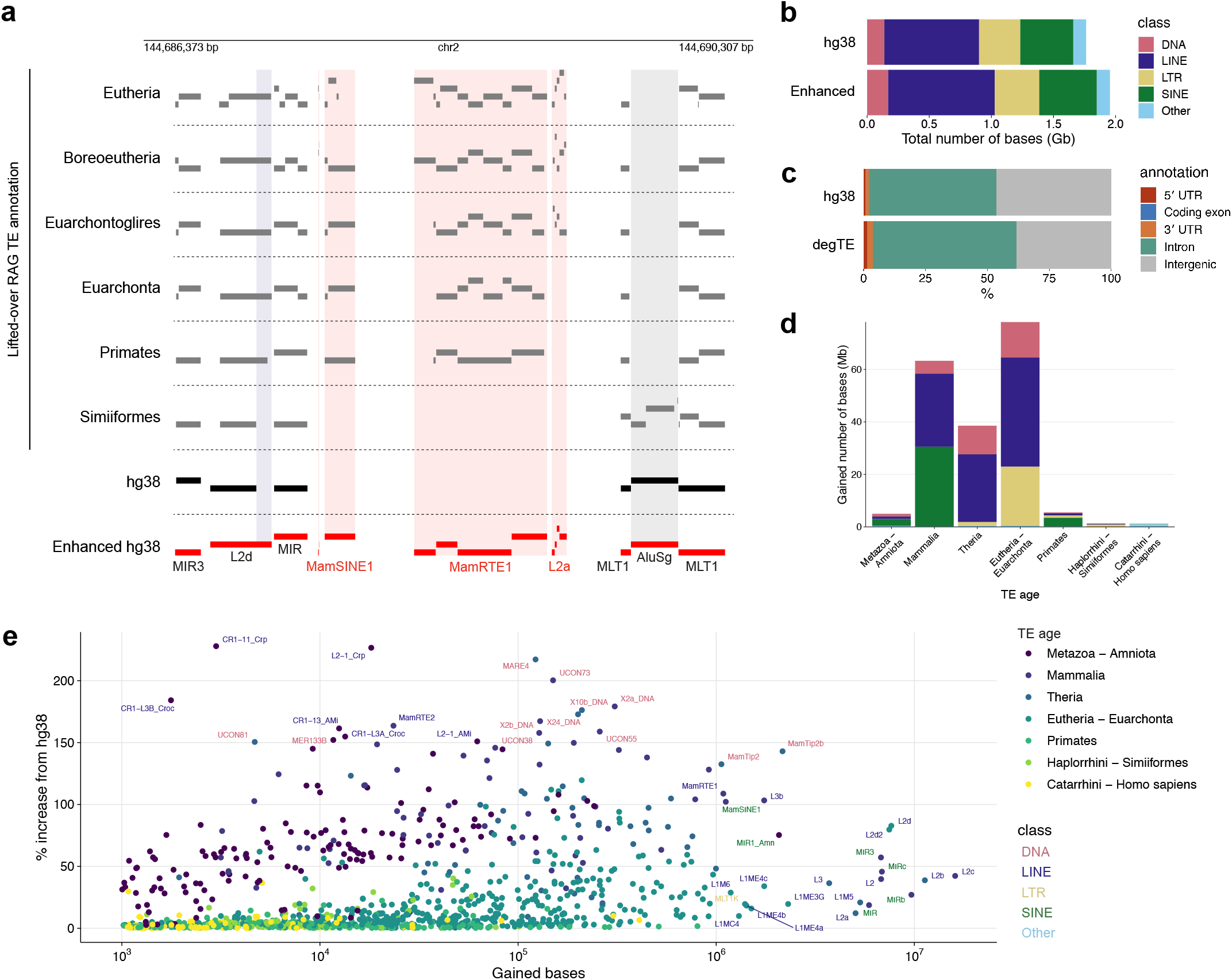
Summary statistics of degTEs. (a) A representative genomic locus where degTEs were identified. Red bars represent enhanced TE annotation produced by merging hg38 TE (black bars) and lifted-over RAG TE (grey bars) annotations. Newly identified integrants are highlighted in red shades, and those extending existing hg38 TEs are shown in blue shades. The grey shade represents an integrant belonging to a primate-specific subfamily, AluSg. (b) TE class distributions in the hg38 and enhanced TE annotations. (c) Proportions of hg38 TEs and degTEs overlapping with distinct genomic annotations. (d) Evolutionary age distribution of degTEs. (e) For the TE subfamilies that gained more than 1 kb by the enhanced annotation, the number of bases gained and percent increase relative to the hg38 annotation are plotted with a colour representing the TE age. The subfamilies that gained either more than 1 Mb or 150% are labelled with their subfamily name in a colour of a corresponding TE class.

The enhanced annotation, a union of degTEs and hg38 TEs, increased TE coverage in the human genome by 10.8% without major changes in the proportions of TE classes (Fig. 3b). The vast majority of degTEs were found in either intronic or intergenic regions (Fig. 3c), similarly to TEs already annotated in hg38. They are distributed across all chromosomes (Supplementary Fig. 1c), with a weak but significant bias (*P* < 2.2e-16) towards regions harbouring fewer TEs already annotated in hg38 (Supplementary Fig. 1d).

Ninety-seven percent of the degTEs are derived from TE subfamilies that emerged sometime between the mammalian and Eutherian common ancestors (Fig. 3d, Supplementary table 1). Especially, the L2 and MIR subfamilies that emerged during this evolutionary time window significantly contribute to the new annotation, each gaining approximately an additional 10 Mb (Fig. 3e). Noteworthy, several ancient DNA transposon subfamilies increased by more than 150% in the enhanced annotation relative to the hg38 TE annotation.

Next, we tested how much homology degTE sequences retained, compared with the corresponding RAG inserts. For this, we performed a sequence homology search on degTEs found as novel integrants against the same TE database used for the TE annotation without the inclusion threshold imposed in RepeatMasker. This analysis revealed that more than 20% of the degTEs displayed significant homology to the same subfamily as the one found for the corresponding sequences in the RAGs, whereas the remaining elements were either assigned to different TE subfamily or showed no significant homology to any TEs (Supplementary Fig. 2a). The homology scores obtained for degTEs were expectedly lower than those of the corresponding TEs in the RAGs (Supplementary Fig. 2b). Also, we observed that degTE L2c elements exhibited similar coverage as those already annotated in hg38. The observed 5′ truncation is a typical feature for LINE integrants^15^ (Supplementary Fig. 2c).

Together, these results show that enhanced TE annotation was achieved by recovering evolutionarily old TEs that exhibited higher homology to the consensus in the RAGs without significant difference in the coverage pattern compared to TEs already annotated in hg38. Importantly, while some (∼20%) degTEs may be discovered by simply running RepeatMasker with a lower threshold, the remaining elements will not, presumably because of more extensive mutational changes.

### degTEs contribute regulatory elements and TF binding sites

Given that TE-derived *cis*-regulatory elements (CREs) are prevalent and play crucial gene regulatory roles, we next tested if the discovered degTEs contribute CREs to the human genome. We crossed the enhanced hg38 TE annotation with the genome-wide candidate CRE (cCRE) annotation, which is based on epigenetic signatures in multiple cell types^16^. In addition to the cCREs associated with previously annotated hg38 TEs, we identified 82,474 cCREs associated with degTEs, which correspond to ∼8% of each cCRE category (Fig. 4a).

**Figure 4.**
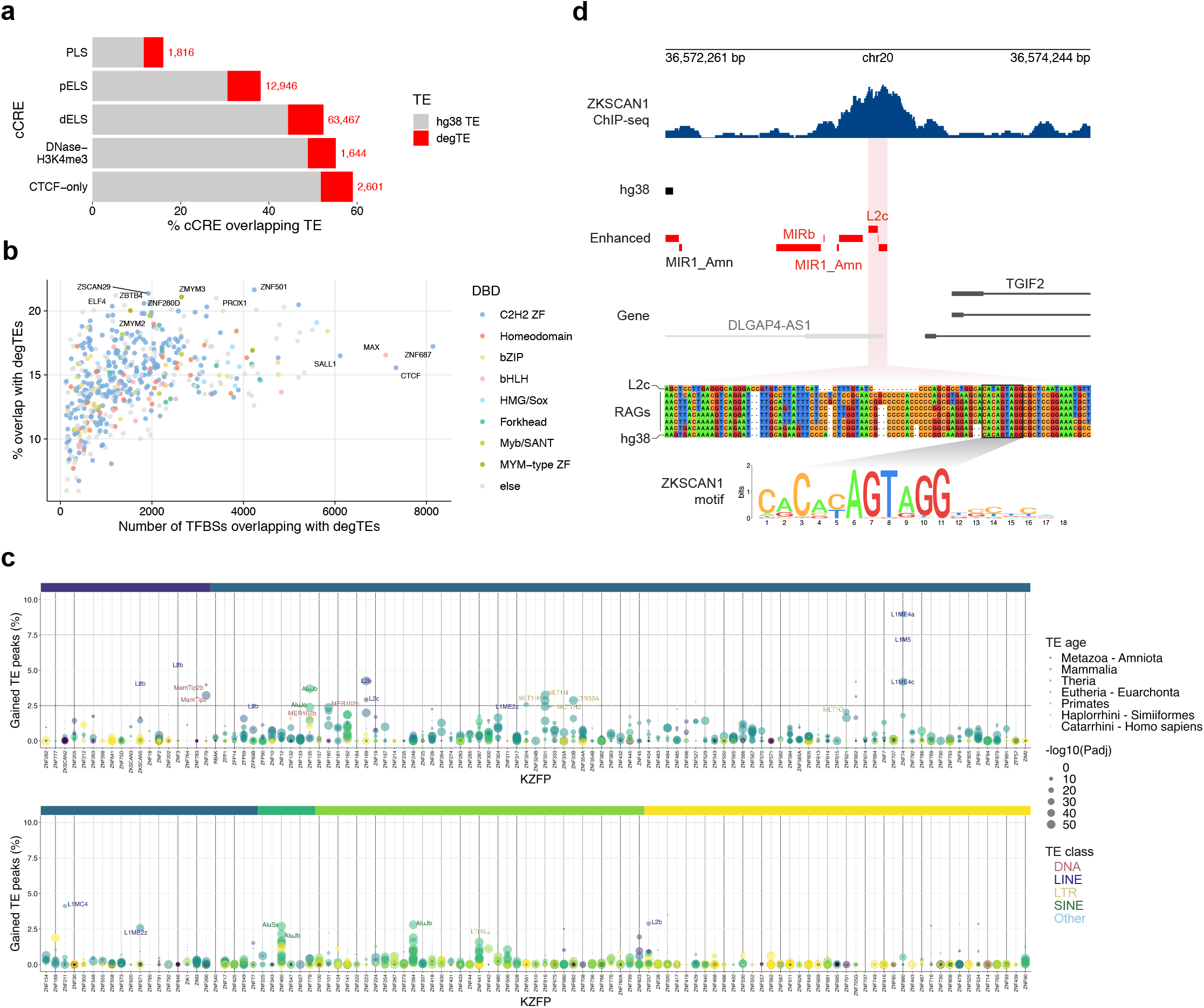
cCREs and TFBSs derived from degTEs. (a) Percentage of cCREs overlapping the hg38 TEs and degTEs. The red numbers indicate cCREs that overlap with one or more degTEs. (b) ENCODE TFBSs overlapping with one or more degTEs. The colours represent DNA-binding domains of the TFs. (c) A summary of degTE-overlapping KZFP binding sites shown in percentage relative to the total number of peaks. Each point represents a TE subfamily and its colour and size indicate its evolutionary age and an adjusted enrichment *P*-value, respectively. The same colour scale is used for the ribbon above to indicate the ages of the KZFPs. (d) A representative genomic region where a degTE-derived KZFP binding site was found. A multiple sequence alignment is shown for the newly found L2c element in the middle of the ZKSCAN1 peak and its corresponding sequences in the RAGs together with the L2c consensus. The sequence matching the ZKSCAN1 binding motif is highlighted.

Next, to see if the degTEs contribute TF binding sites (TFBSs), we crossed the enhanced TE annotation with the ENCODE TFBS data. We identified ∼6,000 degTE-associated TFBSs for each TF (Fig. 4b, Supplementary table 2). Among the TFs with the highest proportion of degTE-derived TFBSs, we found the co-repressor of repressor element-1 silencing transcription (CoREST) complex components, ZMYM2 and its paralogue ZMYM3^17^. ZMYM2 has been reported to be involved in TE silencing^18,19^, which may explain their frequent association with degTEs.

KRAB domain-containing zinc-finger proteins (KZFPs) are major TFs that recognise TEs, usually binding TE subfamilies of a matching evolutionary age^20,21^. We thus tested if degTEs are also bound by KZFPs. We compared degTE annotation with the ChIP-seq peaks from 210 KZFPs displaying significant enrichment for one of the TE subfamilies in the human genome. We found the binding sites of KZFPs that were previously identified as targeting evolutionarily old TE subfamilies to overlap often with degTEs belonging to the same subfamilies (Fig. 4c). Conversely, since only a limited proportion of degTEs are evolutionarily young, KZFPs binding young TEs showed less frequent overlaps with degTEs. We further analysed a KZFP, *ZKSCAN1*, which significantly binds L2 integrants. We found a ZKSCAN1 binding site centred on a degTE belonging to the L2b subfamily (Fig. 4d). A multiple sequence alignment of the degTE and corresponding RAG sequences revealed that the latter were indeed closer to the L2b consensus. However, interestingly, the region corresponding to the ZKSCAN1-binding motif is highly conserved across the RAGs and hg38, suggesting that it has been under purifying selection more than the rest of this TE sequence.

Thus, the recovery of degTEs through the analysis of RAGs revealed the previously unknown association of cCREs and TFBSs with TEs.

### degTEs contribute to chimeric transcripts

Non-canonical chimeric transcripts between TEs and genes, or transpochimeric gene transcripts (TcGTs), have been detected in various cell types including mammalian embryos^6,7^. We thus sought for TcGTs derived from degTEs using previously published RNA-seq data from human embryos^22^, which led us to identify 250 non-canonical exons that involve at least one degTE (Fig. 5a). For instance, a TcGT for *LMOD3* is initiated from a LTR79 degTE located at 30 kb upstream of the first annotated exon for this gene (Fig. 5b). This indicates that some degTEs also contribute to TcGTs by either driving their expression or being included as a non-canonical exon.

**Figure 5.**
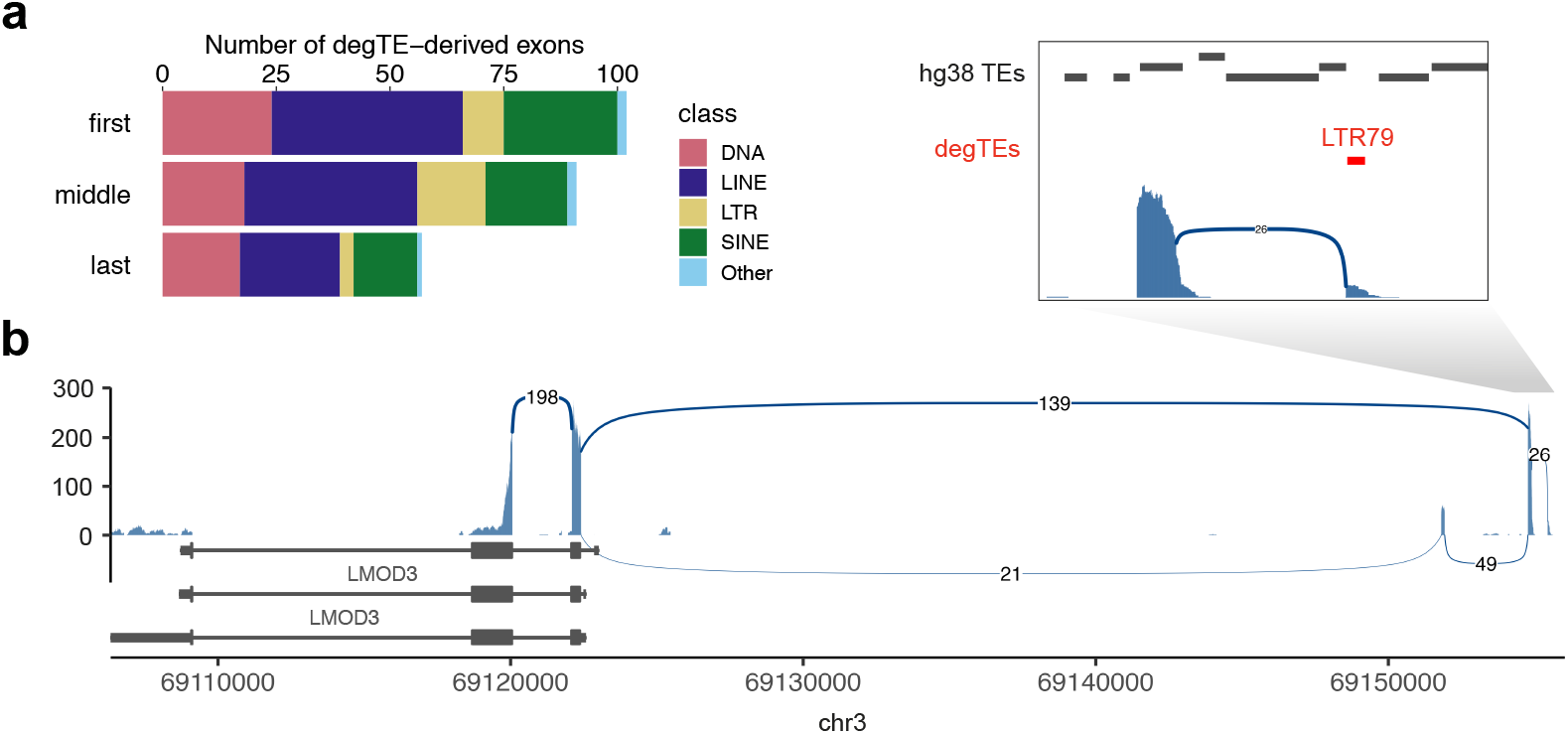
Chimeric transcripts involving degTEs. (a) The numbers of degTE-overlapping exons that contribute to unannotated human embryonic transcripts. (b) A Sashimi plot indicating the number of splicing events observed for a gene where a chimeric transcript with a degTE, LTR79, was found. Splice junctions with more than 10 supporting reads are shown.

## Discussion

Upon discovering transposable elements in the middle of the 20^th^ century, Barbara McClintock coined the term controlling elements for the regulatory potential of TEs^23^, and, two decades later, Eric Davidson and Roy Britten postulated that they played fundamental roles in building regulatory networks^24,25^ Accumulating evidence has now validated this hypothesis, demonstrating that TE-derived proteins, non-coding transcripts, and various *cis*-acting elements are key to evolution of genome regulation and architecture (reviewed in refs.^3,4^).

To study these TE co-option events, TE annotation plays a central role. TE annotation can be either repository-based or performed *de novo* (reviewed in ref.^26^). The former most commonly uses a software called RepeatMasker, which annotates TEs by looking for sequences that resemble those catalogued in a reference repeat database such as Repbase or Dfam. Thus, inherently, this method can only detect TEs with a reasonably high similarity to the consensus sequences. Also, since sequences are annotated based solely on a given genome, mutations gained over evolution are not taken into consideration, which may result in false classifications. On the other hand, *de novo* annotation is generally considered to be more robust to sequence variations from the consensus, and thus is more likely to discover TEs than repository-based annotation. However, it is also prone to high false-positive rates. A report estimates that P-clouds^27^, which predicted the highest number of elements in the human genome, may entail a false-positivity rate (FPR) of ∼60%, which is much higher than the estimated 0.4% FPR of RepeatMasker^28^.

In this study, we describe a novel method to obtain a more complete TE annotation by incorporating information from multiple RAGs, while still utilising a commonly used repository-based annotation method. This allows for not only more sensitive recovery of degenerate TEs, but also more interpretable results of newly annotated TEs, since corresponding sequences in RAGs are available with nucleotide-resolution alignment to consensus sequences. Our method may also be useful in updating already-annotated TE integrants based on the annotation of corresponding sequences in RAGs, since these are closer to the ancestral state of the integrants.

Our method relies on two major factors: reconstruction of ancestral genomes and TE annotation with RepeatMasker. For the former, although the ancestral genomes used were reconstructed from up to 241 genomes, the result still is only a prediction of the ancestral states, which might involve a certain degree of errors and might affect the TE prediction accuracy at specific genomic loci. For the latter, any TE annotation software applies a threshold to call elements. Although RepeatMasker is no different and may report some false positives, the major benefit of our method is that the TEs in the species of interest (SOI) were also called with the same method and threshold. In other words, TEs found in RAGs and SOI are called with the same confidence.

Ancestral reconstructions of several TE subfamilies have been achieved using multiple integrants either within or across species^29–31^. Campitelli et al. took TE reconstruction to the next level by employing ancestral genome reconstruction and achieved the reconstruction of the full-length LINE-1 progenitor sequence^32^. However, to our knowledge, our study is the first attempt where whole reconstructed ancestral genomes were used to mine degenerate TEs in a genome of interest. Our approach should be applicable to different species or TE annotation methods, although high-quality RAGs would be critical in achieving accurate degTE annotation.

The largest increase in the enhanced annotation was seen for the L2 subfamilies, each gaining approximately 10 Mb of degTEs corresponding to ∼90% increase from the current hg38 annotation (Fig. 3e). Multiple studies have reported an enrichment of L2 elements in various CREs, including promoters, enhancers, and DNA loop anchors^33,34^. Also, L2 elements have been reported to be a source of microRNAs and their target sites^35^. Thus, newly annotated L2 elements in our enhanced annotation will help achieve a full grasp of the L2-mediated gene regulatory system.

Our approach identified novel cCREs, TFBSs, and TcGTs associated with old TEs, which had not been found with current hg38 TE annotation, further reinforcing the validity of Britten and Davidson’s original hypothesis on the role played by mobile genetic elements in the genome-wide dissemination of response-mediating sequences. In addition, our enhanced TE annotation method as well as the resulting improved human TE annotation would open up wide-ranged applications. For example, since this approach achieves locus-level reconstruction of each element, it will facilitate defining how individual TE subfamilies have propagated across genomes and how various TE-derived elements have evolved over time. With the ever-increasing number of sequenced genomes, RAGs will become more accurate and older RAGs will be reconstructed, which will further extend the breadth of applications of this method.

## Methods

### TE annotation

Cactus genomic alignment HAL file was downloaded from the Cactus alignment project website^9^ (https://cglgenomics.ucsc.edu/data/cactus/). All the reconstructed ancestral genomes and the human genome were isolated from the HAL file using the hal2fasta script from HAL tools^14^ (version 2.1) (https://github.com/ComparativeGenomicsToolkit/hal). The corresponding node names of the RAGs used in this study are summarised in Supplementary Table 3. Dfam family database^10^ (Dfam-p1_curatedonly.h5.gz, version 3.6) was downloaded from Dfam website (https://www.dfam.org/) and was used as a reference. Search engine HMMER and species “human” together with the options “-s -a -nolow” were used to run RepeatMasker (ver. 4.1.2-p1) for all the genomes used. RepeatMasker output files were converted to the BED file format for downstream analyses.

TE annotation for each ancestral genome was lifted over to the human genome using halLiftover from HAL tools (version 2.1). For the enrichment analyses shown below, to prevent inflated significant scores, the fragmented annotation after the halLiftover was merged. Specifically, TE annotations that are on the same strand, are within 100 bp from each other, and belong to the same subfamily were merged into a single continuous annotation. For the other statistics, the unmerged fragmented annotation was used.

### TE characterisation

The estimated age of each TE subfamily was obtained from Dfam_curatedonly.embl file downloaded from the Dfam website (https://dfam.org/releases/Dfam_3.6/). In cases where a subfamily is associated with multiple nested clades, the oldest clade was assigned (e.g. “Eutheria; Primates” was reclassified as “Eutheria”).

To annotate genomic loci degTEs fall into, BED files of genic regions, UTRs, introns, and coding exons, were taken from the UCSC Table Browser (http://genome-euro.ucsc.edu/). The intergenic regions were defined as regions outside the genic regions.

### TE and degTE distribution analysis

All the manipulations were done with the R package bedtoolsr^36^ (version 2.30.0-1). The human genome was split into 100-kb bins with bt.makewindows, and the coverage calculation for TE/degTE in each bin were performed by bt.coverage. The hg38 centromere annotation was obtained using the UCSC Table Browser (http://genome-euro.ucsc.edu/).

### degTE homology analysis

A degTE was defined as a genomic region where no TE was annotated in hg38 but was assigned a TE in one of the corresponding loci in the RAGs. The homology of the degTEs to TE sequences was determined by running the nhmmscan function from HMMER software^37^ (version 3.3.2) with the default parameters against the same Dfam database (version 3.6) as in the RepeatMasker annotation.

### L2c alignment to the consensus

The sequences annotated as L2c were obtained with bedtools^38^ getfasta (version 2.30.0) command by running it on hg38 and the Eutherian RAG. 1,000 elements were randomly chosen from each set and were aligned to the consensus sequences obtained from Dfam (version 3.6) with MAFFT^39^ –addfreagments command (version 7.508). The resulting alignment was visualised with Jalview^40^ (version 2.11.2.6).

### degTE overlap with cCRE and TFBS

The human candidate *cis*-regulatory elements (cCREs) data (registry V3) were downloaded from the SCREEN website (https://screen.encodeproject.org/index/cversions). To simplify the categories, the CTCF-bound cCRE categories were merged into other cCRE categories (e.g. both “PLS” and “PLS,CTCF-bound” were collapsed into a single category “PLS”).

Overlaps between hg38 TEs/degTEs and cCREs were defined by “bedtools intersect -f 0.5 - F 0.5 -e” (version 2.30.0), meaning an overlap length must be longer than 50% of either the TE/degTEs or cCRE annotation. With this, we defined cCREs that do not overlap with hg38 TEs but overlap with degTEs.

For the TFBS analysis, we downloaded TF ChIP-seq data generated by the Richard Myers group from the ENCODE website^41^ (https://www.encodeproject.org/). ENCODE dataset ID can be found in the “id” column of Supplementary Table 2. We defined TFBS that do not overlap with hg38 TEs but overlap with degTEs by bedtools using the same parameters used for cCRE as shown above.

### TE enrichment in KZFP binding sites

KZFP binding sites were taken from ref.^21^ Statistical test on the enrichment levels of hg38 TEs in the KZFP binding sites was performed with pyTEnrich software with the default parameters (https://alexdray86.github.io/pyTEnrich/build/html/index.html).

### Chimeric transcripts in human embryos

FASTQ files of RNA-seq on human embryos were obtained from GSE197265^22^. The reads were mapped to hg38 using STAR^42^ (version 2.7.10b) with the following parameters “--outFilterMultimapNmax 20 --alignSJoverhangMin 10 --alignSJDBoverhangMin 1 --outFilterMismatchNmax 999 --outFilterMismatchNoverReadLmax 0.04 --alignIntronMin 20 --alignIntronMax 1000000 --alignMatesGapMax 1000000”. Using the obtained alignment, *de novo* transcriptome assembly was performed with StringTie^43^ (version 2.2.1) with the following parameters “-c 1 -f 0.01 -p 1 -j 1 -a 10”. Of the resulting transcripts, exons that were identified in all the biological replicates for each embryonic stage were kept for downstream analyses. The filtered exon annotation obtained from the *de-novo* identified transcripts were then crossed with degTEs that do not overlap with the canonical exons.

To visualise the splicing event of the RNA-seq data, ggsashimi^44^ was used.

## Data availability

Associated data including the RepeatMasker output of all the RAGs as well as their corresponding loci in hg38 were deposited on Zenode (doi:10.5281/zenodo.7716409).

## Acknowledgements

We thank the Trono lab members for constructive discussions. Most computational works were conducted on the high-performance computing cluster developed and maintained by Scientific IT and Application Support (SCITAS) at EPFL. This work was supported by grants from the European Research Council (KRABnKAP, No. 268721; Transpos-X, No. 694658) and the Swiss National Science Foundation (310030_152879 and 310030B_173337) to D.T.; and EMBO Postdoctoral Fellowship (ALTF 1287-2020) and JSPS Overseas Research Fellowship to W.M.

## Author contributions

W.M. conceived and designed the study, performed the analyses, and wrote the manuscript; E.P. supported the analyses and reviewed the manuscript; D.T. reviewed and edited the manuscript, and provided expertise and feedback.

## Figure legends

**Supplementary Figure 1.**
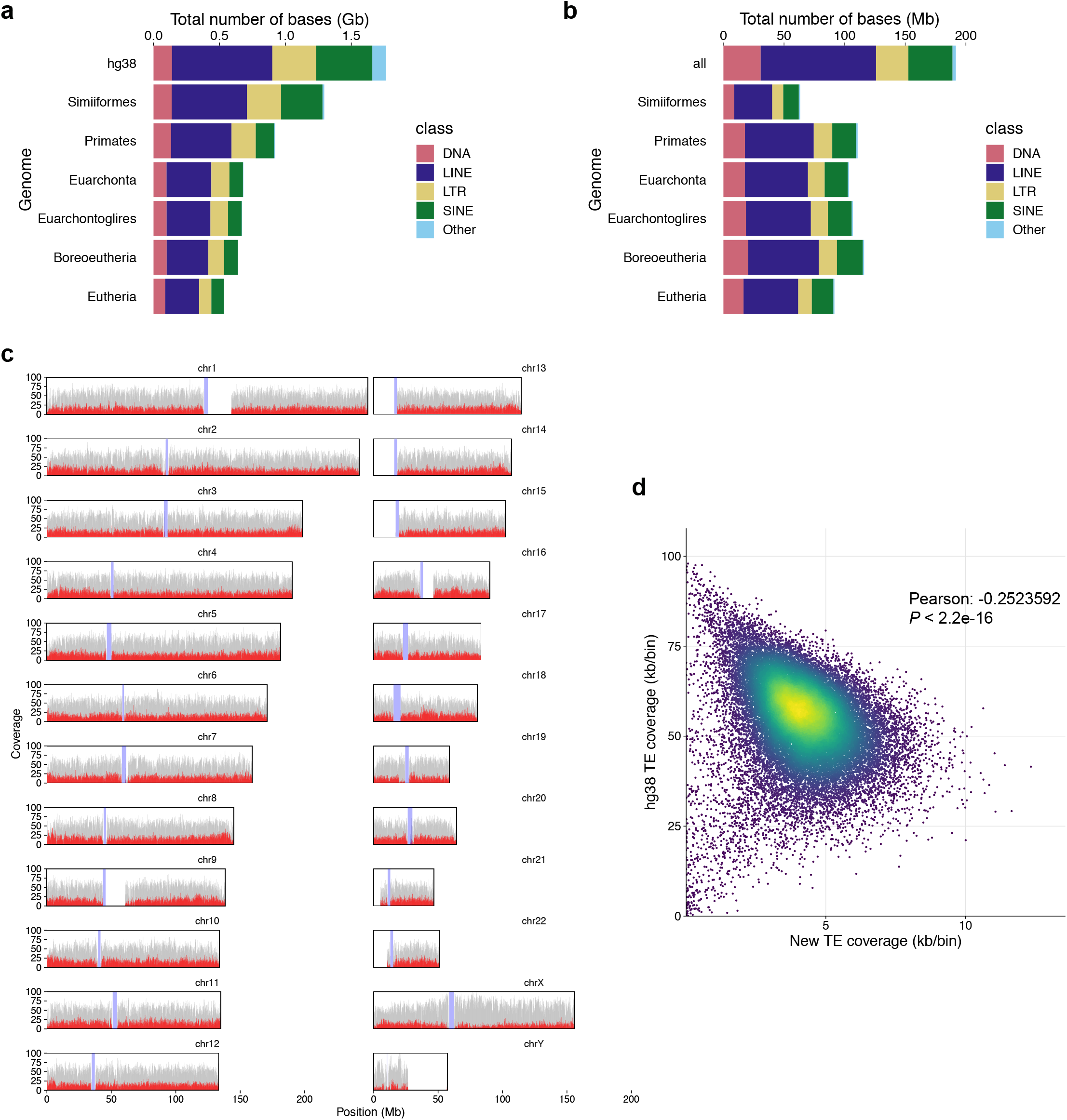
Continued summary statistics of degTEs related to Figure 3. (a) The number of TE bases of in the hg38 TEs and that lifted over to hg38 from each RAG. (b) Coverage of the lifted-over TEs from each RAG that do not overlap with the existing hg38 TE annotation. “all” represents the number combining the lifted-over annotations from all the RAGs. (c) Genome-wide distribution of the hg38 TE (grey) and degTE (red) annotations. The y-axis represents a coverage (kb/bin) for 100-kb bins. The values for the degTEs were multiplied by four for a visualisation purpose. The blue shades represent telomeric regions. (d) The same coverage data as (c) are shown in a scatter plot. Pearson’s correlation coefficient and *P*-value are shown.

**Supplementary Figure 2.**
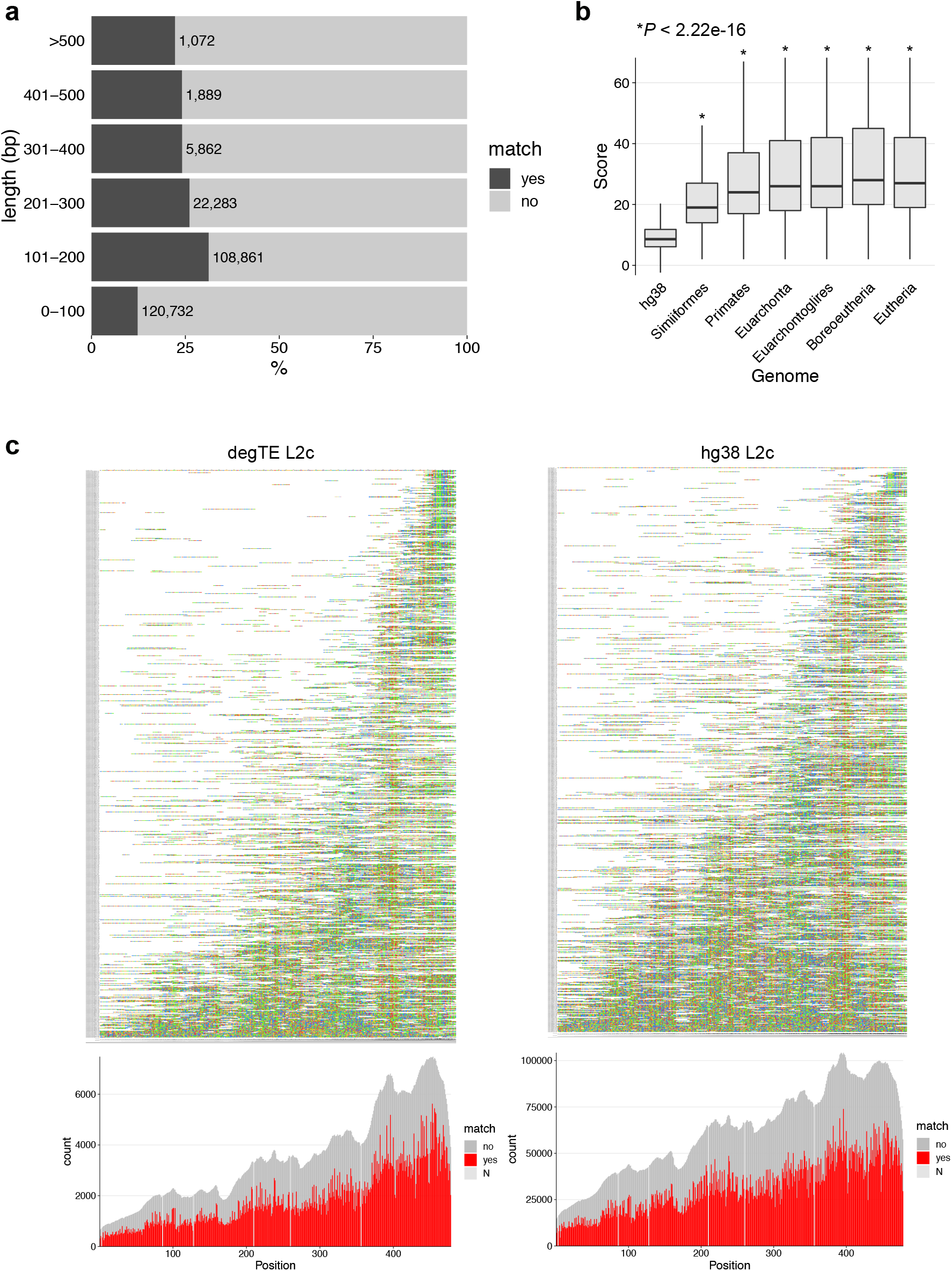
Homology scores and discovery bias observed for degTEs. (a) degTEs corresponding to novel integrants were divided into bins for distinct lengths, and the percentages and numbers of elements are highlighted for the ones exhibiting significant homology to the same TE subfamilies as the ones found in the corresponding regions in the RAGs. (b) Bit score distribution of degTEs and their corresponding sequences in the RAGs. *P*-values from Mann-Whitney *U* test comparing the distributions of degTEs with TEs in each RAG are shown. (c) Annotated L2c integrants aligned to the consensus. The left panes show L2c integrants in the Eutherian RAG that contribute degTEs, and the right panes show those annotated in hg38. The alignment plots were made based on randomly chosen 1,000 elements from each genome. Below, coverage plots over the L2c consensus is shown. For positions where the consensus base is N are shown in light grey.

## References

1. Nurk, S. et al. The complete sequence of a human genome. Science 376, 44–53 (2022).

2. Mills, R. E., Bennett, E. A., Iskow, R. C. & Devine, S. E. Which transposable elements are active in the human genome? Trends Genet. 23, 183–191 (2007).

3. Fueyo, R., Judd, J., Feschotte, C. & Wysocka, J. Roles of transposable elements in the regulation of mammalian transcription. Nat. Rev. Mol. Cell Biol. 1–17 (2022).

4. Jangam, D., Feschotte, C. & Betrán, E. Transposable Element Domestication As an Adaptation to Evolutionary Conflicts. Trends Genet. 33, 817–831 (2017).

5. Modzelewski, A. J., Gan Chong, J., Wang, T. & He, L. Mammalian genome innovation through transposon domestication. Nat. Cell Biol. 24, 1332–1340 (2022).

6. Peaston, A. E. et al. Retrotransposons regulate host genes in mouse oocytes and preimplantation embryos. Dev. Cell 7, 597–606 (2004).

7. Göke, J. et al. Dynamic Transcription of Distinct Classes of Endogenous Retroviral Elements Marks Specific Populations of Early Human Embryonic Cells. Cell Stem Cell 16, 135–141 (2015).

8. Smit, A. F. A., Hubley, R. & Green, P. 2013--2015. RepeatMasker Open-4.0. Preprint at (2021).

9. Jurka, J. Repbase update: a database and an electronic journal of repetitive elements. Trends Genet. 16, 418–420 (2000).

10. Wheeler, T. J. et al. Dfam: a database of repetitive DNA based on profile hidden Markov models. Nucleic Acids Res. 41, D70–82 (2013).

11. Zoonomia Consortium. A comparative genomics multitool for scientific discovery and conservation. Nature 587, 240–245 (2020).

12. Feng, S. et al. Dense sampling of bird diversity increases power of comparative genomics. Nature 587, 252–257 (2020).

13. Armstrong, J. et al. Progressive Cactus is a multiple-genome aligner for the thousand-genome era. Nature 587, 246–251 (2020).

14. Hickey, G., Paten, B., Earl, D., Zerbino, D. & Haussler, D. HAL: a hierarchical format for storing and analyzing multiple genome alignments. Bioinformatics 29, 1341–1342 (2013).

15. Ostertag, E. M. & Kazazian, H. H., Jr. Biology of mammalian L1 retrotransposons. Annu. Rev. Genet. 35, 501–538 (2001).

16. ENCODE Project Consortium et al. Expanded encyclopaedias of DNA elements in the human and mouse genomes. Nature 583, 699–710 (2020).

17. Hakimi, M.-A., Dong, Y., Lane, W. S., Speicher, D. W. & Shiekhattar, R. A candidate X-linked mental retardation gene is a component of a new family of histone deacetylase-containing complexes. J. Biol. Chem. 278, 7234–7239 (2003).

18. Graham-Paquin, A.-L. et al. ZMYM2 is essential for methylation of germline genes and active transposons in embryonic development. bioRxiv 2022.09.13.507699 (2022) doi:10.1101/2022.09.13.507699.

19. Owen, D., Aguilar-Martinez, E., Ji, Z., Li, Y. & Sharrocks, A. D. ZMYM2 controls transposable element transcription through distinct co-regulatory complexes. bioRxiv 2023.01.24.525372 (2023) doi:10.1101/2023.01.24.525372.

20. Wolf, D. & Goff, S. P. Embryonic stem cells use ZFP809 to silence retroviral DNAs. Nature 458, 1201–1204 (2009).

21. Imbeault, M., Helleboid, P.-Y. & Trono, D. KRAB zinc-finger proteins contribute to the evolution of gene regulatory networks. Nature 543, 550–554 (2017).

22. Zou, Z. et al. Translatome and transcriptome co-profiling reveals a role of TPRXs in human zygotic genome activation. Science 378, abo7923 (2022).

23. Mcclintock, B. Controlling elements and the gene. Cold Spring Harb. Symp. Quant. Biol. 21, 197–216 (1956).

24. Britten, R. J. & Davidson, E. H. Gene regulation for higher cells: a theory. Science 165, 349–357 (1969).

25. Davidson, E. H. & Britten, R. J. Regulation of gene expression: possible role of repetitive sequences. Science 204, 1052–1059 (1979).

26. Goerner-Potvin, P. & Bourque, G. Computational tools to unmask transposable elements. Nat. Rev. Genet. (2018) doi:10.1038/s41576-018-0050-x.

27. de Koning, A. P. J., Gu, W., Castoe, T. A., Batzer, M. A. & Pollock, D. D. Repetitive elements may comprise over two-thirds of the human genome. PLoS Genet. 7, e1002384 (2011).

28. Caballero, J., Smit, A. F. A., Hood, L. & Glusman, G. Realistic artificial DNA sequences as negative controls for computational genomics. Nucleic Acids Res. 42, e99 (2014).

29. Ivics, Z., Hackett, P. B., Plasterk, R. H. & Izsvák, Z. Molecular reconstruction of Sleeping Beauty, a Tc1-like transposon from fish, and its transposition in human cells. Cell 91, 501–510 (1997).

30. Miskey, C., Izsvák, Z., Plasterk, R. H. & Ivics, Z. The Frog Prince: a reconstructed transposon from Rana pipiens with high transpositional activity in vertebrate cells. Nucleic Acids Res. 31, 6873–6881 (2003).

31. Dewannieux, M. et al. Identification of an infectious progenitor for the multiple-copy HERV-K human endogenous retroelements. Genome Res. 16, 1548–1556 (2006).

32. Campitelli, L. F. et al. Reconstruction of full-length LINE-1 progenitors from ancestral genomes. Genetics (2022) doi:10.1093/genetics/iyac074.

33. Cao, Y. et al. Widespread roles of enhancer-like transposable elements in cell identity and long-range genomic interactions. Genome Res. 29, 40–52 (2019).

34. Roller, M. et al. LINE retrotransposons characterize mammalian tissue-specific and evolutionarily dynamic regulatory regions. Genome Biol. 22, 62 (2021).

35. Petri, R. et al. LINE-2 transposable elements are a source of functional human microRNAs and target sites. PLoS Genet. 15, e1008036 (2019).

36. Patwardhan, M. N., Wenger, C. D., Davis, E. S. & Phanstiel, D. H. Bedtoolsr: An R package for genomic data analysis and manipulation. J Open Source Softw 4, (2019).

37. Wheeler, T. J. & Eddy, S. R. nhmmer: DNA homology search with profile HMMs. Bioinformatics 29, 2487–2489 (2013).

38. Quinlan, A. R. & Hall, I. M. BEDTools: a flexible suite of utilities for comparing genomic features. Bioinformatics 26, 841–842 (2010).

39. Katoh, K., Misawa, K., Kuma, K.-I. & Miyata, T. MAFFT: a novel method for rapid multiple sequence alignment based on fast Fourier transform. Nucleic Acids Res. 30, 3059–3066 (2002).

40. Waterhouse, A. M., Procter, J. B., Martin, D. M. A., Clamp, M. & Barton, G. J. Jalview Version 2--a multiple sequence alignment editor and analysis workbench. Bioinformatics 25, 1189–1191 (2009).

41. Luo, Y. et al. New developments on the Encyclopedia of DNA Elements (ENCODE) data portal. Nucleic Acids Res. 48, D882–D889 (2020).

42. Dobin, A. et al. STAR: ultrafast universal RNA-seq aligner. Bioinformatics 29, 15–21 (2013).

43. Pertea, M. et al. StringTie enables improved reconstruction of a transcriptome from RNA-seq reads. Nat. Biotechnol. 33, 290–295 (2015).

44. Garrido-Martín, D., Palumbo, E., Guigó, R. & Breschi, A. ggsashimi: Sashimi plot revised for browser- and annotation-independent splicing visualization. PLoS Comput. Biol. 14, e1006360 (2018).

