## Supplementary table 3 for "Ancestral genome reconstruction enhances transposable element annotation by identifying degenerate integrants"

| Node | Name |
|---|---|
| fullTreeAnc239 | Eutheria |
| fullTreeAnc238 | Boreoeutheria |
| fullTreeAnc115 | Euarchontoglires |
| fullTreeAnc114 | Euarchonta |
| fullTreeAnc110 | Primates |
| fullTreeAnc110point5 | Simiiformes |
